# LncRNA Miat and Metadhedrin maintain a treatment resistant stem-like niche of medulloblastoma cells

**DOI:** 10.1101/2021.09.23.461558

**Authors:** Kai-Lin Peng, Harish N. Vasudevan, Dennis T. Lockney, Rachel Baum, Ronald C. Hendrickson, David R. Raleigh, Adam M. Schmitt

**Affiliations:** Division of Translational Oncology, Department of Radiation Oncology, Memorial Sloan Kettering Cancer Center, New York, NY, USA; Departments of Radiation Oncology and Neurological Surgery, University of California San Francisco, San Francisco, CA, USA; Microchemistry and Proteomics, Memorial Sloan Kettering Cancer Center, New York, NY, USA

## Abstract

Medulloblastoma is a pediatric brain tumor arising from the cerebellum and brainstem that is curable with surgery and aggressive cytotoxic therapy including craniospinal irradiation and chemotherapy. Preclinical models indicate that a small pool of de-differentiated, stem cell-like medulloblastoma cells are resistant to cytotoxic treatment and contribute to medulloblastoma relapse after aggressive therapy. Here, we identify *Miat* as a Shh and Myc regulated long noncoding RNA (lncRNA) that is required for maintenance of a treatment-resistant medulloblastoma stem-like phenotype. Loss of *Miat* delays medulloblastoma formation in genetically defined mouse models of Shh medulloblastoma and enforces differentiation of tumorigenic stem-like medulloblastoma cells into a non-tumorigenic cell state. Miat facilitates treatment resistance in part by downregulating p53 signaling and impairing radiation induced cell death in stem-like MB cells, which can be reversed by therapeutic inhibition of Miat with antisense oligonucleotides. The RNA binding protein Metadhedrin (Mtdh), which is associated with resistance to cytotoxic therapy in numerous types of cancer, co-localizes with Miat in stem-like medulloblastoma cells. Further, loss of Mtdh activates p53 signaling and reduces tumorigenicity in stem-like medulloblastoma cells. Taken together, these data reveal a critical role for the lncRNA Miat in sustaining a treatment resistant pool of tumorigenic stem-like medulloblastoma cells.

## Introduction

Medulloblastoma (MB) is the most common malignant brain tumor in children (1). Treatment with high dose chemotherapy and craniospinal irradiation can cure medulloblastoma, but frequently at the expense of neurocognitive toxicities (2). While tumor classifications based on molecular profiling have revealed pathogenic drivers of the disease and differentiate prognostic groups, attempts to dose de-escalate craniospinal irradiation has been hindered by increased risks of treatment failure, even in the Wnt group of MB patients who generally have the best prognosis (3, 4). Stem-like cancer cells have been postulated to be a reservoir of treatment resistance in medulloblastoma and other brain tumors (5-8). Identifying and targeting mechanisms that maintain treatment resistant stem-like medulloblastoma cells could provide an opportunity to reduce the risk of treatment failure and ultimately support strategies to de-escalate the curative cytotoxic regimen.

The Sonic Hedgehog (Shh) group of medulloblastoma accounts for the largest proportion of patients with the disease. Outcomes in Shh medulloblastoma are highly variable and include favorable and unfavorable subtypes. Indeed, Shh medulloblastomas with *TP53* mutations account for some of the worst prognoses, perhaps due to deficiencies in p53-dependent cell death following cytotoxic therapy (9, 10). p53 signaling also plays an important role in preventing cell dedifferentiation and downregulation of p53 is a requirement for the generation of induced pluripotent stem cells and stem-like cancer cells (11, 12). Furthermore, subpopulations of stem-like medulloblastoma cells have been associated with treatment resistance (7, 8, 13). Therefore, understanding the mechanisms the facilitated the maintenance of stem-like medulloblastoma and their treatment resistance could improve treatment outcomes in the disease.

Long noncoding RNAs (lncRNAs) are increasingly recognized as central actors in and cancer phenotypes (14). Prior work in our lab and others have demonstrated that lncRNAs regulate p53 signaling and cellular differentiation (15, 16). Since many lncRNAs mediate mechanisms that are highly tissue selective or context specific, they are attractive targets for drug development since toxicity and off-target effects are anticipated to be minimal (17, 18). Furthermore, advances in antisense drug development have allowed for efficient targeting and degradation of RNAs, including lncRNAs, by systemic or CNS restricted drug delivery (19, 20). Identification of therapeutically targetable lncRNAs involved in the maintenance of medulloblastoma cells with a stem-like phenotype could augment conventional therapy, improve tumor control, and potentially allow for de-escalation of cytotoxic therapy to reduce neurocognitive toxicity.

## Results and Discussion

To identify lncRNA candidates that could participate in Shh medulloblastoma differentiation, we annotated lncRNAs that were differentially expressed in Shh medulloblastoma from *Math1-Cre*, SmoM2^C^ mice, which are frequently used to model the disease. Here, the expression of a constitutively active, oncogenic point mutation of *Smo* (*SmoM2*^*c*^) is restricted to cerebellar granule neuron progenitors (CGNP) using the conditional transgene *Math1-Cre*. RNA sequencing of *Math1-Cre*; *SmoM2*^*c*^ MB revealed that 642 annotated lncRNAs were differentially expressed in tumors compared to age matched normal cerebella (**Figure 1A, Supplemental Figure 1A**)(21). We examined whether lncRNAs differentially expressed in medulloblastoma were regulated during the differentiation of cerebellar granule neuron progenitors (CGNPs), the likely cell of origin of Shh medulloblastoma(22-24). Of the 642 lncRNAs differentially expressed in medulloblastoma, we identified 113 lncRNAs that changed dynamically during CGNP differentiation and cerebellar development, 44 of which with increased expression in medulloblastoma; and 70 with reduced expression in medulloblastoma compared to normal developing cerebellum (**Supplemental Figure 1B**).

**Figure 1.**
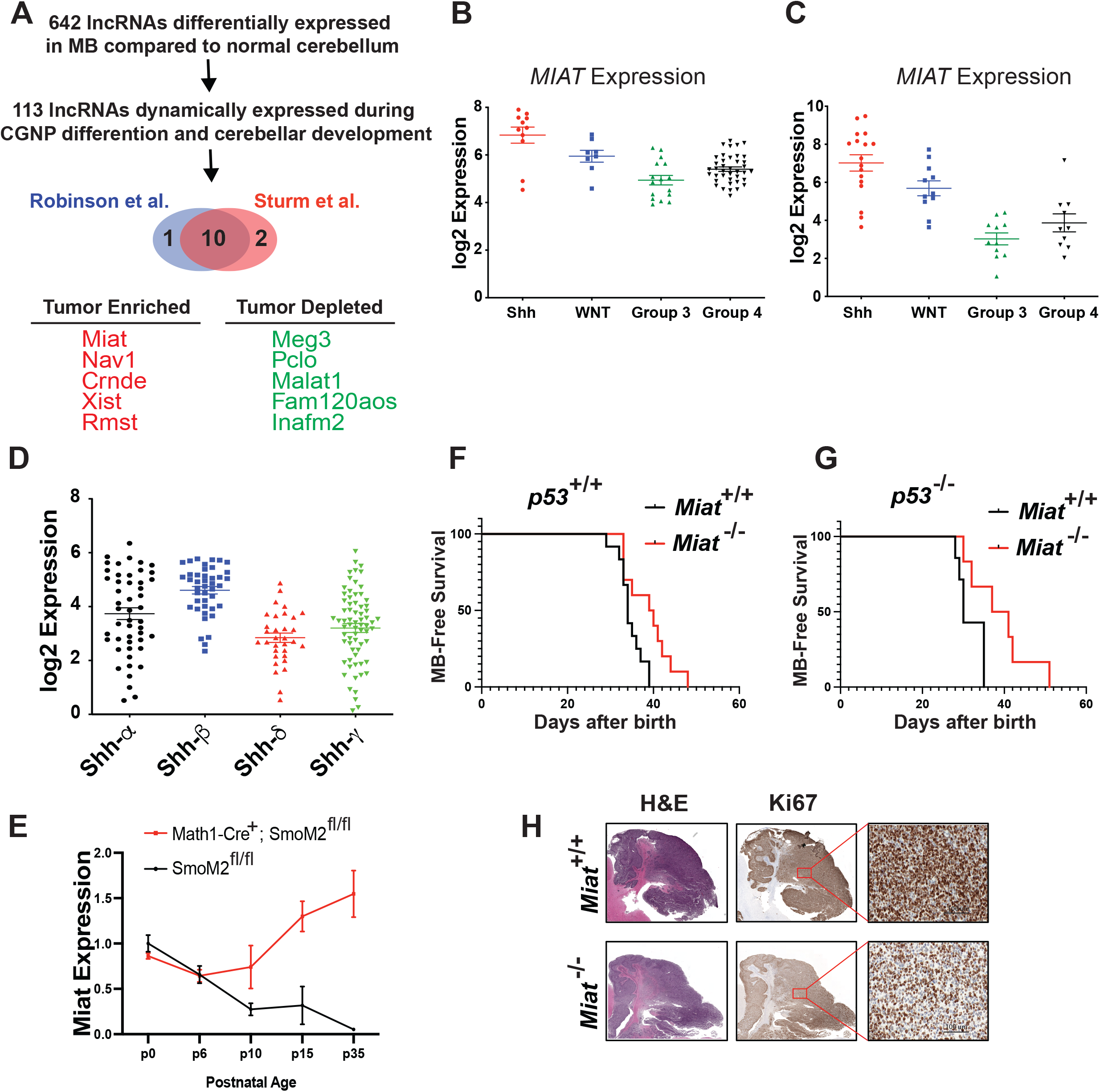
*Miat* is a developmentally regulated, Shh associated lncRNA that enhances MB development. **A**, Flow chart illustrating criteria to identify candidate lncRNAs involved in Shh MB differentiation. **B-C**, *MIAT* expression in molecular groups in Robinson et al. (**B**) and Sturm et al. (**C**). **D**, *MIAT* expression by subgroups of Shh MB. **E**, *Miat* expression in the cerebella of mice of Math1-cre+, SmoM2^fl/fl^ (red line) and cre-, SmoM2^fl/fl^ mice (black line) from postnatal day 0 through 35. **F**, Kaplan-Meier survival curves in Math1-cre+, SmoM2^fl/fl^ Shh MB mice in the *p53*^*+/+*^ background, n=12 *Miat*^*+/+*^ with median survival of 34 days, n=10 *Miat*^*-/-*^ with median survival of 39.5 days, *: p <0.05, log-rank test. **G**, Kaplan-Meier survival curves in Math1-cre+, SmoM2^C^ Shh MB mice in the *p53*^*-/-*^ background, n=7 *Miat*^*+/+*^ with median survival of 30 days, and n=6 *Miat*^*-/-*^ mice with a median survival of 38.5 days, ** p<0.01, log-rank test. **H**, Representative H&E and Ki67 stains of midline sagittal sections through the cerebella from of Math1-Cre; SmoM2^fl/fl^ *Miat*^*+/+*^ and *Miat*^*-/-*^ mice.

To further enrich for lncRNAs that contribute to Shh medulloblastoma, we filtered lncRNAs differentially expressed in the Shh group of human medulloblastoma compared to other groups. Using expression data from previously published human MB datasets(25, 26), we identified 10 medulloblastoma-associated lncRNAs that change dynamically during CGNP differentiation and for which expression differed in the Shh group MB compared to other groups (Figure 1A). Among these lncRNAs, *MIAT* was of particular interest since *MIAT* expression was enriched in human Shh MB (**Figure 1B-C**). Moreover, a Shh-specific enhancer element was noted 12kb upstream of the *MIAT* transcription start site (**Supplemental Figure 1C**), *MIAT* is expressed most highly in the Shh-β subgroup of high risk infant Shh medulloblastoma (**Figure 1D**) (27, 28), and enriched during early cerebellum development at the earliest stages of CGNP neurogenesis and repressed at later stages of development following CGN differentiation (**Figure 1E**). To determine whether *Miat* contributes to Shh MB development, we crossed *Math1-Cre*; *SmoM2*^*c*^ with *Miat*^*-/-*^ mice(29) in both *p53*^*+/+*^ and *p53*^*-/-*^ backgrounds since p53 mutations are strongly associated with aggressive Shh MB subtype with an exceptionally poor prognosis (30). Homozygous genetic loss of *Miat* prolonged median survival in both *p53*^*+/+*^ and *p53*^*-/-*^ MB animals, indicating Miat supports MB development in multiple contexts of Shh MB (**Figure 1F-G**). Homozygous genetic deletion of *Miat* also reduced the proliferative fraction of cells as measured by Ki67 staining and partially restored cerebellar architecture (**Figure 1H**), indicating *Miat* supports the growth of Shh MB.

Since *Miat* is highly expressed in the earliest stages of CGNP development, we next examined whether Miat regulates the differentiation of MB cells. To study the role of Miat in the differentiation of Shh MB, we isolated stem-like MB cells from *Math1-Cre*; *SmoM2*^*c*^ MB by *in vitro* propagation in defined serum-free neural stem cell medium (31, 32). A stem-like phenotype in these cells was confirmed by differentiation (**Supplemental Figure 2A**) and confirmation of tumorigenic potential upon engraftment in the posterior fossa of mice (data not shown). In stem-like MB cells, treatment with smoothened agonist (SAG) induced the expression of *Miat* while Shh signaling antagonist cyclopamine downregulated *Miat* expression, indicating the Shh signaling regulates the expression of *Miat* (**Supplemental Figure 2B**). To understand how *Miat* is regulated in response to Shh signaling, we performed knockdown of Gli2 and n-Myc, transcription factors in the Shh pathway, as well as c-Myc which is required for pluripotency and maintenance of stem-like cancer cells and examined the effect on *Miat* expression. Knockdown of Gli2 modestly decreased the expression of *Miat*, while knockdown of n-Myc or c-Myc substantially downregulated *Miat* expression (**Supplemental Figure 2C-D**). In human DAOY cells, a Shh MB cell line, anti-myc ChIP recovered a region of the *MIAT* promoter that is also associated with histone H3K27ac defined enhancer element upstream of *MIAT* that is specific to Shh group medulloblastoma(33), indicating that the Shh group specific enhancer is a myc bound enhancer (**Supplemental Figures 1C, 2E**). These data indicate that *Miat* is a myc-regulated lncRNA in the Shh pathway.

To elucidate the function of Miat, we generated stem-like MB cells with *Miat* knockdown using CRISPR inhibition (CRISPR-i). qRT-PCR confirmed efficient knockdown of *Miat* by multiple different sgRNAs targeting the *Miat* TSS (**Figure 2A**). RNA-sequencing revealed that *Miat* knockdown resulted in substantial changes in genes involved in stem cell differentiation, apoptosis, neural development, cell-substrate adhesion, and regulation of smoothened signaling (**Figure 2B**) and notably with significant enrichment of canonical p53-dependent genes amongst genes more highly expressed in the *Miat* knockdown samples (FDR adjusted P<10^−4^ by KEGG pathway analysis). As stem-like cancer cells and impair p53-dependent induced cell death have been linked to resistance to cytotoxic therapies, we were interested to examine the implications of p53 dysregulation by Miat on MB responses to cytotoxic therapy. qRT-PCR confirmed that knockdown of *Miat* was associated with increased expression of p53-regulated genes involved in apoptosis and cell cycle arrest, *Puma, Bax*, and *Cdkn1a*. which are necessary for efficient killing of medulloblastoma cells by cytotoxic therapies such as radiation (**Figure 2C**), suggesting that *Miat* expressing stem-like MB cells may be relatively resistant to cell death by cytotoxic therapy. Consistent with a role for Miat in resistance to cytotoxic therapy, knockdown of *Miat* in stem-like MB cells significantly increased the fraction of dead stem-like MB cells following radiation therapy and lowered the threshold for cytotoxicity in stem-like MB cells following radiation therapy (**Figures 2D-E, Supplemental Figure 2F**). In contrast, knockdown of *Miat* in Med1 cells, a differentiated mouse Shh MB cell line that expressed *Miat*, had no effect on sensitivity to radiation, suggesting that Miat’s role in promoting tumor initiation and treatment resistance is limited to the stem-like MB cells. (**Supplemental Figures 2G-H**). Thus, Miat downregulates the expression of p53 pathway effector genes in stem-like MB cells and maintains a treatment resistant stem-like MB cell phenotype.

**Figure 2.**
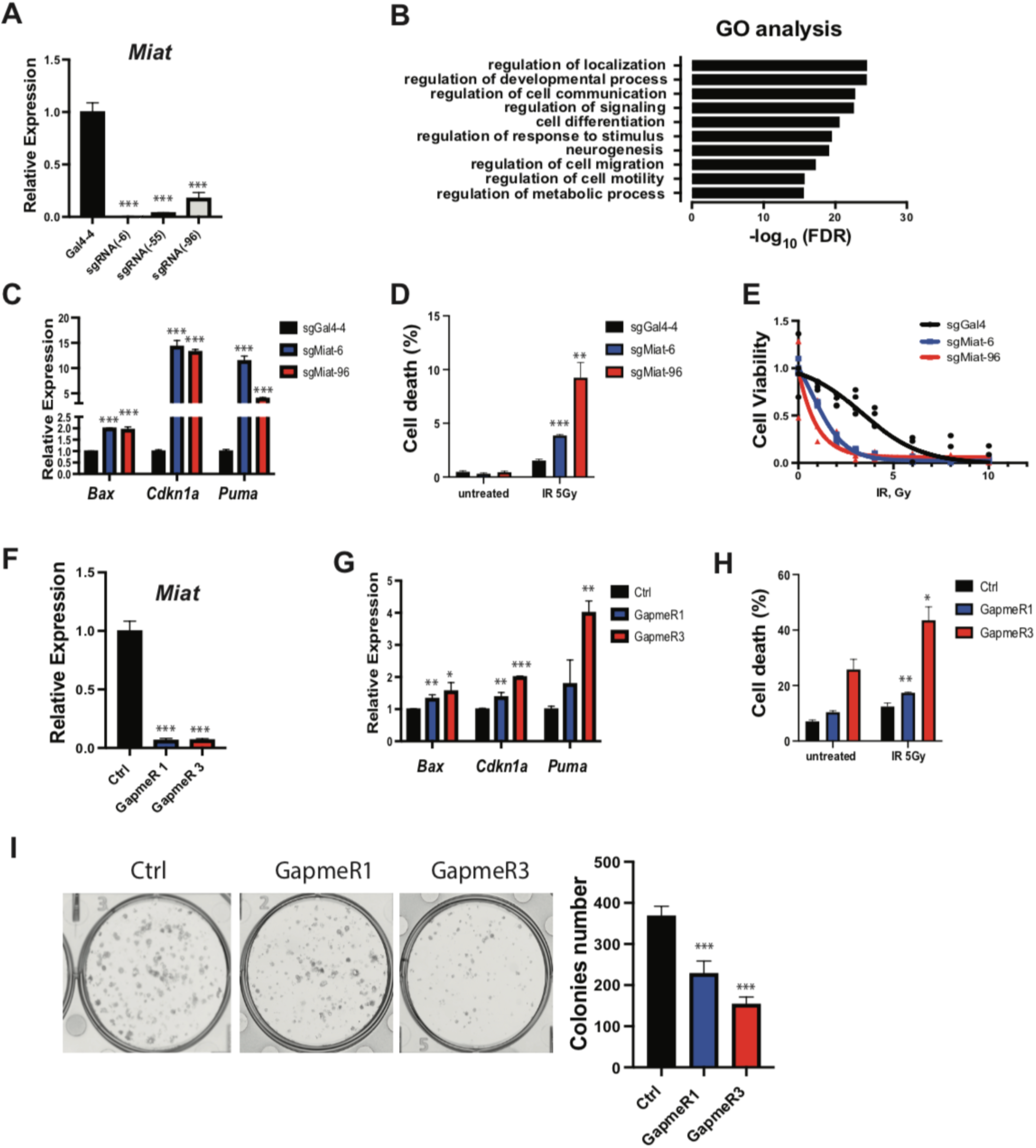
Miat sustains a treatment-resistant stem-like medulloblastoma niche and is therapeutically targetable. **A**, qRT-PCR of *Miat* expression in stem-like MB cells with 3 sgRNAs targeting CRISPRi to *Miat* or control nontargeting sgRNA (Gal4-4). **B**, Gene ontology (GO) terms associated with genes differentially expressed in Miat knockdown stem-lime MB cells compared to controls. **C**, Expression of p53-dependent genes *Bax, Cdkn1a* and *Puma* in stem-like MB cells with sgRNAs targeting CRISPRi to *Miat* or control nontargeting sgRNA (Gal4-4) (n=3). **D**, Fraction of dead cells as measured by Propidium iodide (PI) staining 24 hours after irradiation (5 Gy), in stem-like MB cells with sgRNAs targeting CRISPRi to *Miat* or control nontargeting sgRNA (Gal4-4) (n=3). **E**, Cell viability as measured by MTS assay 48 hours after increasing doses of irradiation (IR) (0, 1, 2, 3, 4, 6, 8, 10 Gy), in stem-like MB cells with sgRNAs targeting CRISPRi to *Miat* or control nontargeting sgRNA (Gal4-4) (n=4). **F-G**, Expression of Miat (**F**) and p53-dependent genes *Bax, Cdkn1a* and *Puma* (**G**) in stem-like MB cells treated with ASOs targeting *Miat* or control (n=3). **H**, Fraction of dead cells as measured by Propidium iodide (PI) staining 24 hours after irradiation (5 Gy) in stem-like MB cells treated with ASOs targeting *Miat* or control (n=3). **I**, Colony formation in stem-like MB cells treated with ASOs targeting *Miat* or control, Right: quantification (n=3). *** p-value<0.001; ** p-value<0.01 by Student’s T test.

Nuclear lncRNAs can be efficiently targeted for therapeutic inhibition and *Miat*^*-/-*^ mice are healthy and fertile, making Miat an attractive therapeutic target. To examine whether Miat can be therapeutically targeted, we generated custom Antisense LNA GapmeRs to knockdown Miat in stem-like MB cells. Two GapmeRs achieved efficient knockdown *in vitro*, and induced p53-dependent genes *Puma, Bax*, and *Cdkn1a* similar to knockdown of *Miat* using CRISPRi (**Figure 2F-G**). A single treatment of stem-like MB cells with Miat GapmeRs increased cell death at baseline and following radiation therapy and reduced colony formation *in vitro* (**Figure 2H-I**). Hence, Miat is a candidate therapeutic target in Shh medulloblastoma to reduce the tumorigenicity and treatment resistance of stem-like MB cells.

Our RNA-seq data also demonstrated that knockdown of *Miat* significantly altered the expression of genes associated with cancer stem cells and differentiation. qRT-PCR validated that knockdown of *Miat* reduced the expression of pluripotent cancer stem cell markers such as *Cx3cl1, Cxcr4*, and *Itga6* (**Figure 3A**); and in contrast, upregulated markers or differentiation, like *Gsc, Pax2, Hoxa7, Phf19* (**Figure 3B**). Since *Miat* knockdown in stem-like MB cells was associated with increased expression of differentiation markers and reduced expression of genes associated with the stem-like phenotype, we examined the effect of *Miat* knockdown on *in vitro* colony formation and neurosphere formation, an assay of a stem-like cancer cell phenotype. *Miat* knockdown completely impaired the ability of stem-like MB cells to form colonies and neurospheres (**Figure 3C**) but had only a modest effect on *in vitro* cell proliferation (**Figure 3D**). *Miat* knockdown had no effect on colony formation in Med1-MB cells, a differentiated mouse MB cell line derived from a a *Ptch1*^*+/-*^ mouse (**Supplemental Figure 3A**) (34, 35). Since Miat is required for the maintenance of a stem-like phenotype in MB cells *in vitro*, we next examined whether Miat contributes to medulloblastoma tumor initiation from stem-like cells *in vivo*. Control stem-like MB cells readily form luciferase-expressing medulloblastoma tumors after engraftment in the posterior fossa and subsequently developed rapidly progressive neurologic symptoms including severe ataxia within 8 weeks of engraftment (**Figures 3E-F**). However, in contrast, mice engrafted with *Miat* knockdown stem-like MB cells showed no evidence of medulloblastoma formation by luciferase and had prolonged survival compared to mice engrafted with control stem-like MB cells (**Figures 3E-F**). H&E and Ki67 staining demonstrated that control stem-like MB cells formed tumors in posterior fossa that were uniformly positive for Ki67, however, no tumor cells were found in mice engrafted with stem-like MB cells with *Miat* knockdown (**Figure 3E**). In sum, these data suggest Miat is necessary for medulloblastoma stem-like cell maintenance and tumorigenesis.

**Figure 3.**
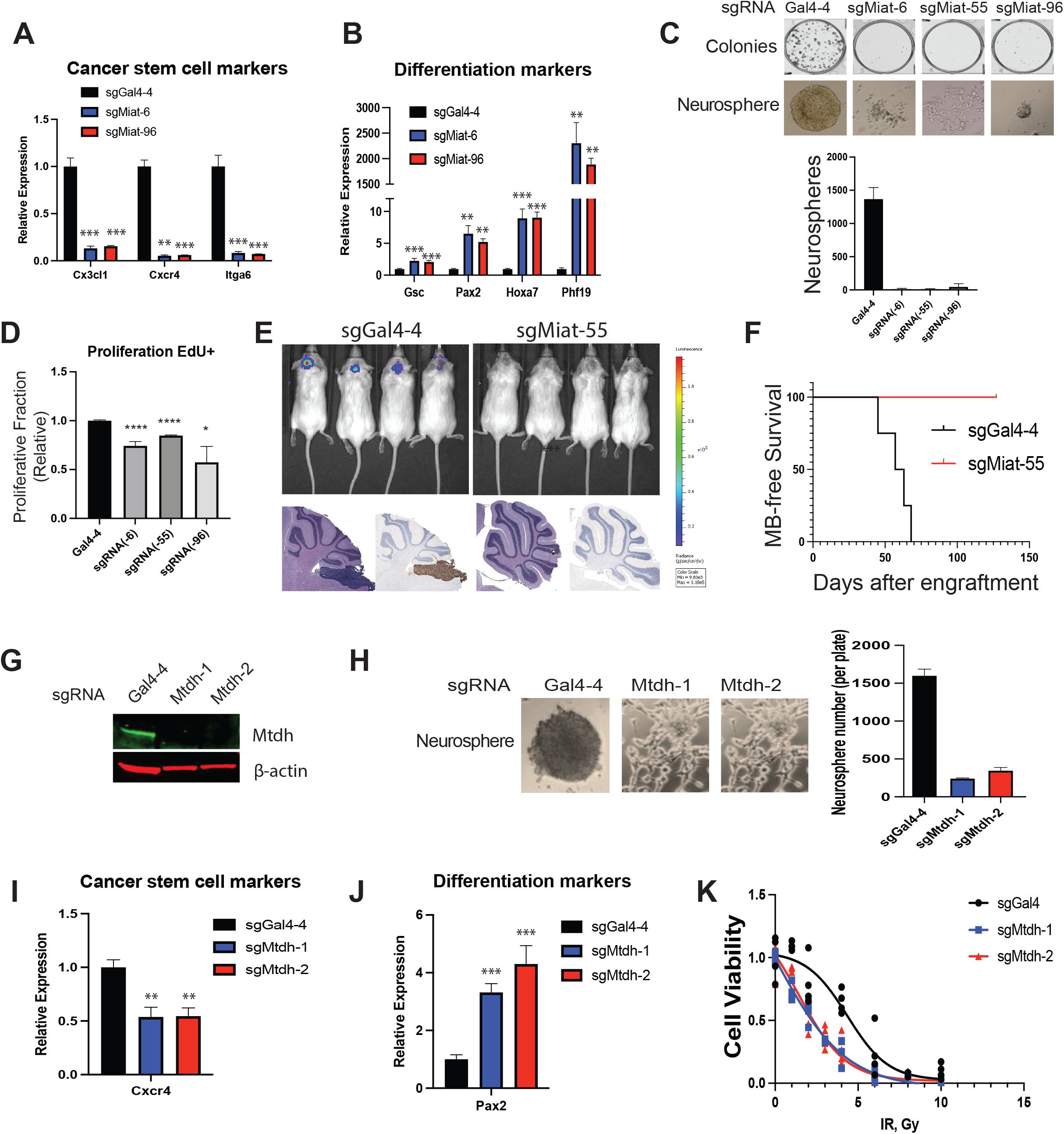
Miat and protein partner Mtdh maintain a stem-like phenotype crucial for tumorigenesis. **A-B**, Expression of genes associated with cancer stem cells (**A**) or stem cell differentiation (**B**) in stem-like MB cells with indicated sgRNAs targeting Miat for CRISPRi or nontargeting control (Gal4-4) (n=3). **C**, Colony formation and neurosphere formation in stem-like MB cells with indicated sgRNAs targeting Miat for CRISPRi or nontargeting control. Knockdown of *Miat* impaired the ability to form colonies (upper panel) and neurospheres (lower panel) (n=3). Below: quantitation of neurosphere formation. **D**, Relative fraction of cells proliferating (EdU+) in stem-like MB cells with indicated sgRNAs targeting Miat for CRISPRi or nontargeting control. **E**, Orthotopic tumor formation of stem-like MB cells with indicated sgRNAs targeting Miat for CRISPRi or nontargeting control with luciferase imaging 2 weeks after engraftment (upper panel) and H&E and Ki67 histopathology at the experimental endpoint (lower). **F**, Kaplan-Meier survival curves for mice orthotopically engrafted with stem-like MB cells with indicated sgRNAs targeting Miat for CRISPRi (n=4) or nontargeting control (n=4). **G**, WB in stem-like MB cells after CRISPR with sgRNAs targeting Mtdh or nontargeting (Gal4-4). Mtdh targeting sgRNAs efficiently eliminated Mtdh protein expression. **H**, Neurosphere formation in stem-like MB cells with *Mtdh*^*+/+*^ (Gal4-4) and *Mtdh*^*-/-*^ (Mtdh-1 and Mtdh-2), right: quantification of neurosphere formation in each condition (n=3). **I-J**, Expression of Miat-regulated, pluripotent cancer stem cell marker *Cxcr4* (**I**) and cellular differentiation marker *Pax2* (**J**) in *Mtdh*^*+/+*^ and *Mtdh*^*-/-*^cells stem-like MB cells (n=3). **K**, Cell viability as measured by MTS assay 48 hours after increasing doses of irradiation (IR) (0, 1, 2, 3, 4, 6, 8, 10 Gy in *Mtdh*^*+/+*^ and *Mtdh*^*-/-*^ stem-like MB cells (n=4 in each group). *** p-value<0.001; ** p-value<0.01; * p-value<0.05 via Student’s T test.

LncRNA mechanisms are frequently mediated by their interactions with other cellular macromolecules such as protein, RNA or DNA. To identify the mechanisms by which Miat functions to maintain the stem-like MB phenotype, we next characterized the proteins that co-localized with Miat in stem-like MB cells. sgRNAs targeting Miat guided the APEX2-dCas13 fusion protein to Miat-enriched nuclear domains and proteins were subsequently labeled by proximity biotinylation (36). Mass spectrometry of recovered biotinylated proteins identified Metadherin (Mtdh), also known as LYRIC, as a highly enriched protein in samples isolated from stem-like MB cells expressing APEX2-dCas13 and sgRNAs targeting Miat (**Supplemental Figure 3B, Supplemental Table 1**). Mtdh is a known RNA binding protein previously shown to contribute to mechanisms of chemoresistance and is required for tumor initiating capacity of cancer cells (37-39). Transient knockdown of Mtdh using siRNAs as well as CRISPR knockout of *Mtdh* (*Mtdh*^*-/-*^) in stem-like MB cells impaired neurosphere formation *in vitro*, similar to knockdown of *Miat* (**Figures 3G-H, Supplemental Figures 3C-F**). Furthermore, gene expression in *Mtdh*^*-/-*^ stem-like MB cells suggested changes associated with differentiation, similar to *Miat* knockdown in stem-like MB cells, with downregulation of stem cell marker Cxcr4 and upregulation of the differentiation marker Pax2 (**Figures 3I-J**). *Mtdh*^*-/-*^ stem-like MB cells, similarly, had increased expression of p53 effector genes *Puma, Bax*, and *Cdkn1a* and reduced stem-like MB cell viability following radiation therapy (**Figure 3K, Supplemental Figures 3G-H**). Thus, Mtdh is necessary for the maintenance of a treatment resistant stem-like population of medulloblastoma cells, phenotypically similar to loss of *Miat*.

These results have identified a new, therapeutically targetable mechanism that maintains a niche of tumorigenic and treatment resistant stem-like MB cells that requires the lncRNA Miat and its interaction with the protein Mtdh. Extensive prior work in preclinical models indicated that Mtdh-dependent processes would be attractive targets for improving cancer sensitivity to cytotoxic therapy (37, 38), but translation to clinic has been hampered by the fact that Mtdh has no ligand binding domain and is considered an undruggable target (40). Further, sterility in *Mtdh*^*-/-*^ males also limits enthusiasm for drug development target Mtdh due to concerns for toxicity. Miat is critical for the maintaining a treatment resistant and tumorigenic potential of stem-like MB cells while simultaneously dispensable for health and normal development, making it an especially attractive therapeutic target in MB. More broadly, these data suggest that conserved cancer mechanisms thought to be undruggable may be vulnerable to modulation by targeting lncRNAs in the pathway.

## Methods

Please refer to Supplemental Methods for details.

## Supporting information

Peng et al Supplemental Text and Figures

## Data Availability

All raw sequencing data will be accessible in the NCBI’s GEO database.

## Study Approval

All animal experiments and procedures were approved by MSKCC’s Institutional Animal Care and Use Committee.

## Author Contributions

HV, DRR and AMS conceived the study. KLP, HNV, DRR, RCH, and AMS designed experiments. DRR and AMS supervised the study. KLP performed all mouse, cellular, and biochemical experiments presented in the figures. HNV and KLP conducted genomics analyses. RCH supervised protein identification by mass spectrometry. DTL and RB performed experiments that contributed importantly to the conception of the study but were not ultimately displayed in the figures. KLP, HV, RCH, DRR, AMS conducted data analyses. The manuscript was prepared by KLP, HNV, DRR, AMS. All authors reviewed the manuscript.

## Acknowledgements

We thank Azusa Tanaka for helpful discussions and comments on the manuscript. Shinichi Nakagawa kindly provided the *Miat*^*-/-*^ mice. Scott Lowe kindly provided the *p53*^*-/-*^ mice. This study was supported by funding from NIH R35GM124909 (to A.M.S.), American Cancer Society RSG-19-158-01-RMC (to A.M.S. and R.C.H.), and the MSKCC Cancer Center Core Grant P30 CA008748.

