## Supplementary material for "LncRNA Miat and Metadhedrin maintain a treatment resistant stem-like niche of medulloblastoma cells": Peng et al Supplemental Text and Figures

### Supplemental Materials

### Detailed Methods

#### Mice

C57BL/6 (#000664), *SmoM2-eYFP*$^{loxP/loxP}$ (#005130), *Math1-Cre* (#011104), Luciferase (#005125), NOD-scid (#005557) mice were purchased from the Jackson Laboratory. *Miat* (*Gomafu*) null Neo-(Acc.No.CDB1377K) mice were acquired from RIKEN (#RBRC10157) (1). p53 null mice were kindly provided by Scott Lowe. Mouse genotyping were performed using Cre, SmoM2, Luc, p53, and Miat primers as indicated. All mice were of the species Mus musculus and maintained on a C57BL/6 background over at least five generations and were handled in compliance with all relevant ethical regulations for animal testing and research as specified by the MSK Institutional Animal Care and Use Committee in approved protocol 15-07-011.

#### Cell Culture

Stem-like MB stem cells were cultured according to established protocols (2). Briefly, tumor-bearing cerebellum was retrieved from MB mice. After dissociation by 50% Accutase (Sigma#A6964-100ML), cells were cultured in 0.1% gelatin coated plates with neurobasal medium (Thermo#21103049), Pen-Strep, Glutamine, N2 (Thermo#17502048), B27 (-RA) (Thermo#12587010), hEGF 25 ng/mL (Thermo#PHG0311), and bFGF 25 ng/mL (Thermo#PHG0311). Med1-MB cells were kindly provided by David R. Raleigh and cultured in DMEM with Pen-Strep, Glutamine. For clonogenic cell survival assays, 1000 of MB cells or 500 of Med1-MB cells were plated per well in triplicate in a 6 well dish and cultured for 9 days. Colonies were fixed with methanol, stained with crystal violet and counted using GelCount™

(Oxford Optronix). Edu incorporation was assayed by Click-iT® Plus EdU Alexa Fluor® 488 Flow Cytometry Assay Kit (Thermo#C10632). Cell viability was measured by MTS assay (Promega#G3580) and propidium iodide (Thermo#P1304MP) either 24 or 48 hr after irradiation (IR) or vismodegib (LC Laboratories#V-4050) treatment.

**Plasmid Construction, Lentivirus, and Retrovirus Production**

Cas9 sgRNAs were cloned into lentiCRISPRv2 (Addgene#52961) using BsmBI site. sgRNA seq: Gal4-4, AACGACTAGTTAGGCGTGTA; *Miat* (-6), AGTAGCCCCTTTGTGAGGCG; *Miat* (-55), CGTTGCTATGGCAGCGCCGC; *Miat* (-96), GATCGCGCCTCCGACCGTC. *Mtdh*-1, CGCCATTGTTCCGCCGGGGG; *Mtdh-2*, AGGCGGCCGCTAAGCGGCGT. APEX2-dCasRx plasmid was purchased from Addgene (#154939). CasRx pre-gRNA cloning backbone (Addgene#109054) was cloned into pBabe-zeo (Addgene#1766) with cutting site NheI. sgRNA cassettes were then cloned into pBabe-zeo-pre-gRNA vector using golden gate cloning with restriction site of BbsI. sgRNA seq of non-target, CACCGGACGGAGGCTAAGCGTC; three oligos of *Miat*, CAGGACTCAAGCAAAGAGATGT, CACTGAACAGAACTGTATGGTG, TGGTGAAATGTGGAAGATTGGC. Lentiviral vectors and retroviral vectors were transfected into LentiX 293T cells (Takara#632180) and Pheonix-293T, respectively, and supernatants were concentrated using Lenti-X or Retro-X concentrator prior to use.

**Antisense Oligo (ASO), siRNA and Compound Administration**

Stem-like MB cells were nucleofected with either 250 nM of ASOs or 100 nM of siRNAs using Mouse Neuron Nucleofector Kit (Lonza#VVPG-1001) according to the manufacturer's instructions. 48 hr after nucleofection, cells were harvested for RT-qPCR or Western blot. Seq of

ASO negative control, +A*+A*+C*A*C*G*T*C*T*A*T*A*+C*+G*+C; *Miat* GapmeR1, +T*+A*+G*C*A*C*T*T*T*G*A*T*T*+G*+A*+C; *Miat* GapmeR3, +A*+G*+A*T*G*C*A*G*G*C*G*A*T*+T*+A*+G (+, locked nucleic acids; *, phosphorothioated DNA based). siRNAs were purchased from Thermo silencer select (#4390771) with *Myc* (siRNA ID#s70225), *MycN* (#s70661), *Gli2* (#s66727) and *Mtdh* (#s84566). For Shh agonist and antagonist treatment, stem-like MB cells were treated with either 250 nM of SAG (Sigma#SML1314), or 1 μM of cyclopamine (Stemcell#72072) for 24 hr, and then RNA was extracted for expression analysis. For retinoic acid (RA) treatment, MB was treated with 500 nM RA (Sigma#R2625) for 48 hr.

**Tumor Sphere Assay**

Tumor sphere assay (hanging drop method) was performed as described previously (2). Briefly, 10,000 cells were seeded in the form of 30 μL drops on the inverted lid of the 10 cm petri dish, while the dish contained $Ca^{2+}$ and $Mg^{2+}$ free PBS supplemented with penicillin 100 (U/mL) and streptomycin (100μg/mL). After 48 hr, cells were taken pictures by microscope.

**Mice Allograft Model and Survival Analyses**

For allograft studies, one microliter containing one million of stem-like MB cells was injected into the cerebellum of NOD-scid (JAX#005557) at a position 0 x 2 x 1.5 mm with respect to Lambda by a stereotactic frame. Tumor-bearing mice were monitored daily for health status and movement abnormalities. Tumor luminescence images were acquired by IVIS Spectrum. All mice were euthanized at the onset of symptoms, including 10% weight loss, ataxia, or lethargy. The age of the animal at the time of harvest due to tumor symptoms was recorded. Kaplan-Meier curves were plotted using GraphPad Prism and significance calculated using the Test. The Kaplan–Meier method was used to estimate survival and log-rank tests were used to compare survival between experimental groups.

**Histology and immunohistochemistry**

For histological analysis, mouse brains were removed and fixed with 4% PFA overnight and then transferred to 70% alcohol and embedded in paraffin. Sections were then stained with hematoxylin and eosin or with Ki67 antibody, which were done by core facility-laboratory of comparative pathology at MSKCC.

**Quantitative RT-PCR and Chromatin immunoprecipitation (ChIP)**

qRT-PCR and ChIP were performed as described previously (3). Primer sequences are listed in Table1. Significance calculated by t-test, error bars show standard error of the mean.

**Western blotting and antibodies**

The commercial Abs were listed as below: Mtdh (Abcam#ab124789), Myc (Abcam#39688), Myc (Santa-Cruz#sc53993), β-actin (Abcam#6276), β-tubulin (Genetex#GTX101279), H3K27ace (Abcam#4729), mouse IgG (Sigma#12-371) and rabbit IgG (CST#2729S).

**APEX labelling and Streptavidin Bead Enrichment of Biotinylated Proteins.**

APEX2 labeling was performed as described previously (4). Briefly, 24 hr prior to labeling, stem-like MB cells stably expressing APEX2-dCasRx fusion, transposase and non-target gRNA or sgMiat were treated with doxycycline (400 ng/mL). Then, biotin-phenol (final concentration 500 μM) was directly added into culture medium for 30 min at 37°C. Later, $H_2O_2$ was added to media to a final concentration of 1mM to induce biotinylation. After very gently swirling for 1 min, the media was decanted as quickly as possible and the cells were washed three times with PBS containing 100 mM sodium azide, 100 mM sodium ascorbate, and 50 mM TROLOX (6-hydroxy-2,5,7,8-tetramethylchroman-2-carboxylic acid). The cell pellets then were lysed with RIPA lysis buffer (50 mM Tris, 150 mM NaCl, 0.1% sodium dodecyl sulfate (SDS), 0.5% sodium deoxycholate, 1% Triton X-100, protease inhibitor) followed by streptavidin bead enrichment. Biotinylated proteins were then eluted by boiling the beads in 50 μL of 4× protein loading buffer supplemented with 20mM dithiothreitol (DTT) and 2 mM biotin and ran on SDS-polyacrylamide gel electrophoresis (PAGE) gel. SDS-PAGE gel then was stained by silver staining. Differential bands were undergone LC-MS/MS to identify protein ID. Protein ID was listed in Table 2.

**RNA seq and analysis**

Quality control of FASTQ files was performed with FASTQC, and after trimming of adapter sequences, reads were further filtered to remove bases that did not have an average quality score of 20 within a sliding window across 4 bases (http://www.bioinformatics.babraham.ac.uk/projects/fastqc/). Reads were subsequently mapped to the mouse reference genome mm10 using HISAT2 with default parameters. Transcript abundance estimation in transcripts per million (TPM) and differential expression analysis were performed using DESeq2 (5). We identified differentially expressed transcripts with an adjusted p-value < 0.1 and further filtered significant genes based on an expression cutoff (TPM > 1) and fold change threshold (| log2FC | > 1).

**Supplemental Figure 1. *MIAT* is Ah group-specific lncRNA involved in MB formation.**

**A**, Volcano plot generated by RNA sequencing of the medulloblastomas from 3 P35 Math1-$Cre^{+}$; $SmoM2^{fl/fl}$ mice relative to the cerebella of 3 P35 $SmoM2^{fl/fl}$ littermate controls. 642 genes are differentially expressed with a false discovery rate (FDR) <0.05, differentially expressed lncRNAs in both tumor and control groups, applying p-value < 0.0001 as threshold (red dots). LncRNAs of particular interest are identified. FC, fold change. **B**, Differentially expressed lncRNAs during normal mouse cerebellum development. Cerebellum RNA-seq data analyzed from (6). **C,** H3K27ace ChIP-seq reveals an enhancer located ~12kb upstream of *MIAT* transcription start site (TSS) specific to Shh subtype MB. ChIP-seq data analyzed from (7).

**Supplemental Figure 2. *Miat* is targeted by Shh signaling and regulates cytotoxic responses to radiation therapy exclusively in stem-like MB cells. A**, Expression of stem cell markers in stem-like MB cells after treatment with 500 nM retinoic acid for 48 hours to induce differentiation in comparison to untreated controls. **B**, Gene expression as measured by qRT-PCR of indicated Shh pathway genes in stem-like MB cells treated with Shh agonist SAG, Shh signaling antagonist cyclopamine, or vehicle only as a control (n=3). **C**, Expression of selected genes in stem-like MB cells after knockdown of transcription factors Gli2, n-Myc and c-Myc*,* as measured by qRT-PCR (n=3). **D**, western blot for c-Myc and n-Myc in stem-like MB cells after siRNA knockdown of n-Myc and c-Myc. **E**, ChIP-qPCR for the enrichment by anti-H3K27ac or anti-c-Myc at two locations within 12kb upstream of the *MIAT* TSS. The -12K site corresponds to the location of the H3K27ac peak noted in **Figure S1C** that is unique to Shh group MB, and is bound by c-Myc. **F,** Cellular morphology and growth of stem-like MB cells with Miat knockdown by CRISPRi or control following 10Gy irradiation or without treatment (n=5). **G,** qRT-PCR for *Miat* expression in Med1-MB cells expressing CasRx and sgRNAs targeting *Miat* (n=3). **H,** Cell viability as measured by MTS 48 hours after irradiation of Med1-MB cells with Miat knockdown or control (n=3). *** p-value<0.001; ** p-value<0.01; * p-value<0.05 via student's t test.

**Supplemental Figure 3. Mtdh co-localizes with *Miat* in stem-like MB cells and is required for neurosphere formation and resistance to radiation therapy. A,** Colony formation in Med1-MB cells with knockdown of Miat or nontargeting control. Right panel showed the quantitative result of colonies (n=3). **B,** Biochemical analysis of biotinylated proteins retrieved from stem-like MB cells stably expressing APEX2-dCas13d and indicate sgRNAs, non-targeted (NT) or targeting *Miat*. Lysates were run on SDS/PAGE and analyzed by silver stain (left) ponceau S (middle) or antibody against Mtdh (right). Red arrow pointed to proteins differentially biotinylated by targeted vs. nontargeted APEX2-dCas13. **C-D**. Expression of *Mtdh* by qRT-PCR (**C**) and western blot (**D**) in stem-like MB cells transfected with siRNAs against *Mtdh* or control siRNAs. **E-F,** Images of neurosphere formation (**E**) and quantification (**F**) in stem-like MB cells after transient *Mtdh* knockdown by siRNAs or treatment with control siRNAs (n=3). **G,** Gene expression by qRT-PCR of p53-dependent genes *Bax*, *Cdkn1a* and *Puma* in $Mtdh^{+/+}$ and $Mtdh^{-/-}$ stem-like MB cells (n=3). **H**, Cellular morphology and growth of stem-like $Mtdh^{+/+}$ and $Mtdh^{-/-}$ MB cells following 10Gy irradiation (n=5). *** p-value<0.001; ** p-value<0.01; * p-value<0.05 via student's t test.

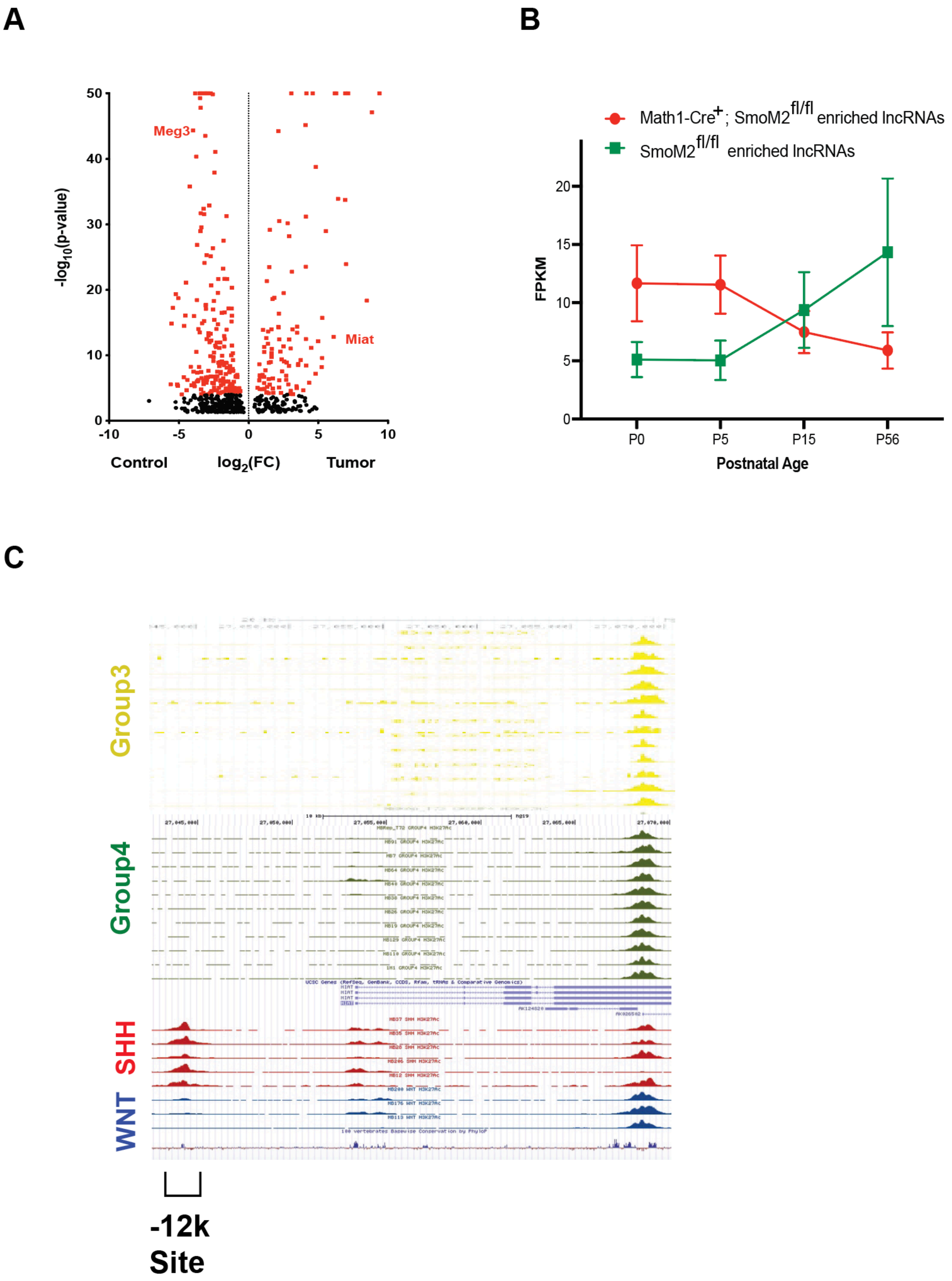
A
B
C
-log10(p-value)
50
40
30
20
10
0
-10
-5
0
5
10
Meg3
Miat
Control
log2(FC)
Tumor
Math1-Cre+; SmoM2fl/fl enriched lncRNAs
SmoM2fl/fl enriched lncRNAs
FPKM
20
15
10
5
0
P0
P5
P15
P56
Postnatal Age
Group3
Group4
SHH
WNT
-12k
Site

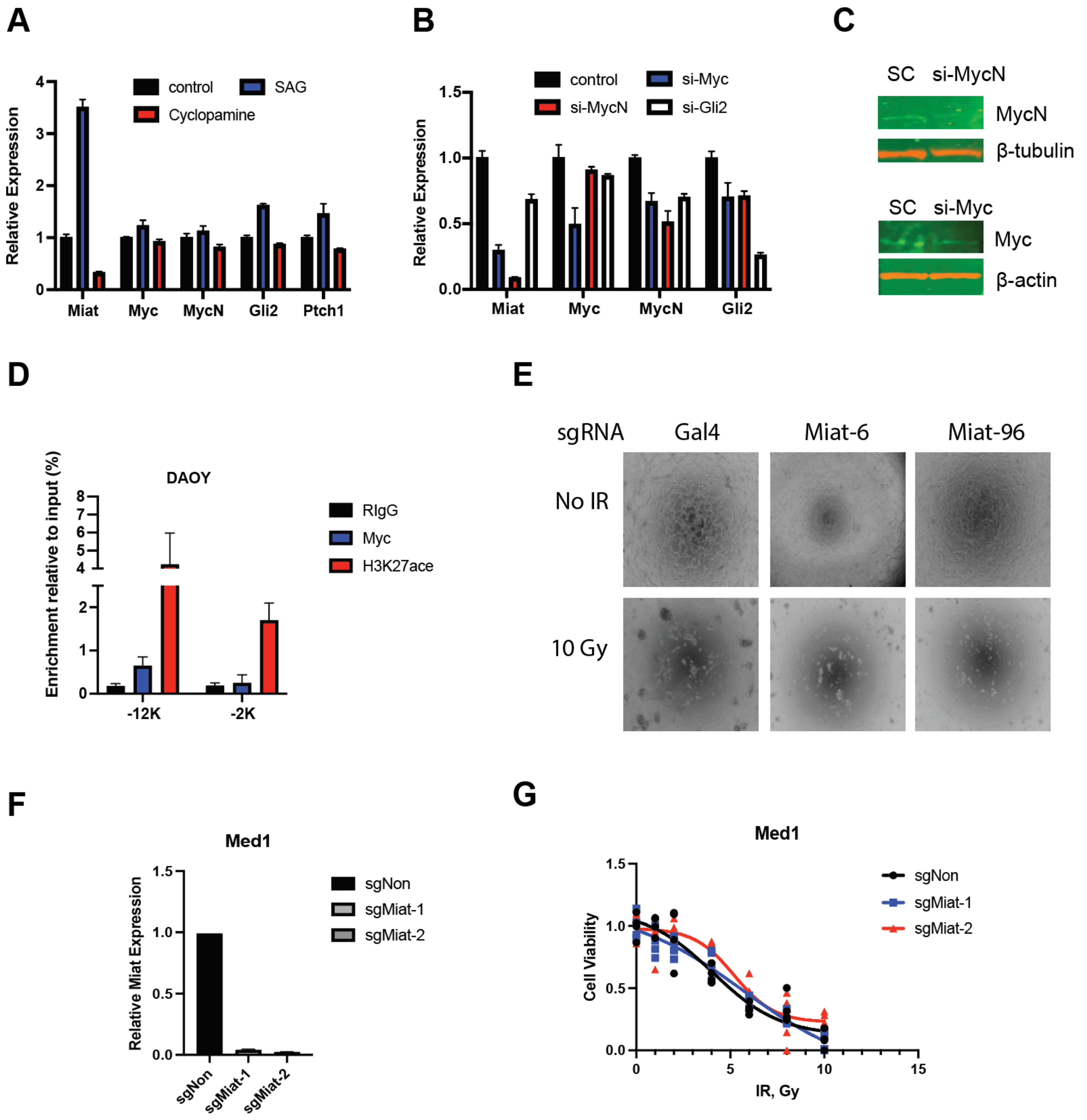
A
control
SAG
Cyclopamine
Relative Expression
Miat
Myc
MycN
Gli2
Ptch1
B
control
si-Myc
si-MycN
si-Gli2
Relative Expression
Miat
Myc
MycN
Gli2
C
SC si-MycN
MycN
β-tubulin
SC si-Myc
Myc
β-actin
D
DAOY
RIgG
Myc
H3K27ace
Enrichment relative to input (%)
-12K
-2K
E
sgRNA
Gal4
Miat-6
Miat-96
No IR
10 Gy
F
Med1
sgNon
sgMiat-1
sgMiat-2
Relative Miat Expression
G
Med1
sgNon
sgMiat-1
sgMiat-2
Cell Viability
IR, Gy

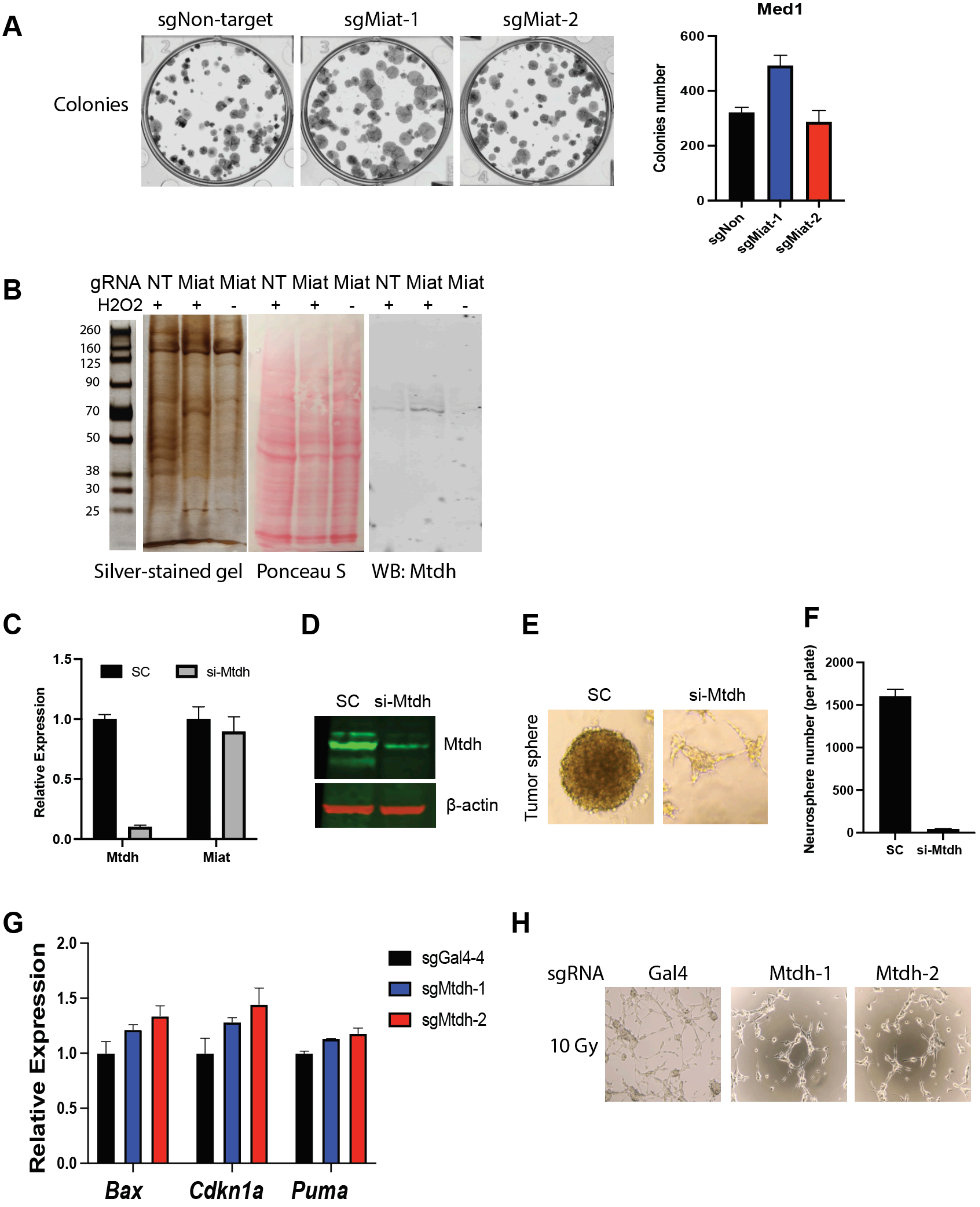

A
sgNon-target
sgMiat-1
sgMiat-2
Colonies
Med1
Colonies number
600
400
200
0
sgNon
sgMiat-1
sgMiat-2
B
gRNA NT Miat Miat NT Miat Miat NT Miat Miat
H2O2 + + - + + - + + -
260
160
125
90
70
50
38
30
25
Silver-stained gel
Ponceau S
WB: Mtdh
C
1.5
1.0
0.5
0.0
Relative Expression
SC
si-Mtdh
Mtdh
Miat
D
SC si-Mtdh
Mtdh
β-actin
E
SC
si-Mtdh
Tumor sphere
F
Neurosphere number (per plate)
2000
1500
1000
500
0
SC
si-Mtdh
G
Relative Expression
2.0
1.5
1.0
0.5
0.0
sgGal4-4
sgMtdh-1
sgMtdh-2
Bax
Cdkn1a
Puma
H
sgRNA
Gal4
Mtdh-1
Mtdh-2
10 Gy
